# Age-dependent remodeling of the sciatic proteome in 5xFAD mice can be attenuated by exercise or donepezil treatment to maintain neuromuscular function

**DOI:** 10.1101/2025.07.01.662603

**Authors:** Matthew H. Brisendine, Dijanira Q. Nieves-Esparcia, Orion S. Willoughby, John R. Brown, Daniel S. Braxton, Shelby N. Henry, Collin McCoin, John P. Thyfault, Jill K. Morris, Steven Poelzing, Robert W. Grange, Charles P. Najt, Joshua C. Drake

## Abstract

Background: Alzheimer’s disease (AD) progresses along a continuum for years to possibly decades prior to cognitive decline and clinical diagnosis. Preclinical AD is associated with neuromuscular dysfunction. We previously characterized early neuromuscular impairment prior to cognitive decline at 4 months of age in the 5xFAD mouse model of AD. However, the underlying cause(s) for peripheral nerve dysfunction leading to impaired skeletal muscle torque production are not understood, therefore limiting interventional capacity. We hypothesized that either voluntary wheel running or donepezil treatment, begun prior to neuromuscular decline, would delay manifestation of neuromuscular impairment in 5xFAD mice.

Methods: Sciatic nerves from 5xFAD and wild-type (WT) mice were analyzed by tandem mass tag (TMT)-labeled proteomics at 3, 4, and 7 months to investigate proteome remodeling. Separate cohorts, using 3-month-old 5xFAD mice and WT littermates given voluntary wheel access for 4 weeks or treated with the acetylcholinesterase inhibitor donepezil to test if neuromuscular dysfunction could be attenuated. Afterwards, we assessed tibial nerve stimulated plantar flexion torque and sciatic nerve compound (motor) neuron action potential (CNAP) in-vivo at 4 months. Additionally, we performed TMT-labeled proteomics to ascertain the effect of exercise and donepezil treatments on sciatic proteome.

Results: Sciatic nerves in 5xFAD mice exhibited proteomic remodeling from 3 to 4 months, particularly in pathways linked to mitochondrial turnover, calcium handling, lipid metabolism, and inflammation, coinciding with onset of neuromuscular dysfunction. Both exercise and donepezil attenuated in nerve-stimulated muscle torque and CNAP dysfunction. Both exercise and donepezil attenuated proteomic remodeling of the sciatic nerve involving mitochondrial-centric processes through both shared and independent mechanisms.

Conclusions: Declines in neuromuscular function may be pre-clinical identifiers for AD that share pathway similarities with noted central effects of the pathology on the brain. Our findings highlight the importance of a systemic approach to AD pathology and importance of disease state in interventional efficacy.

**Graphical abstract.:** Created in Biorender.

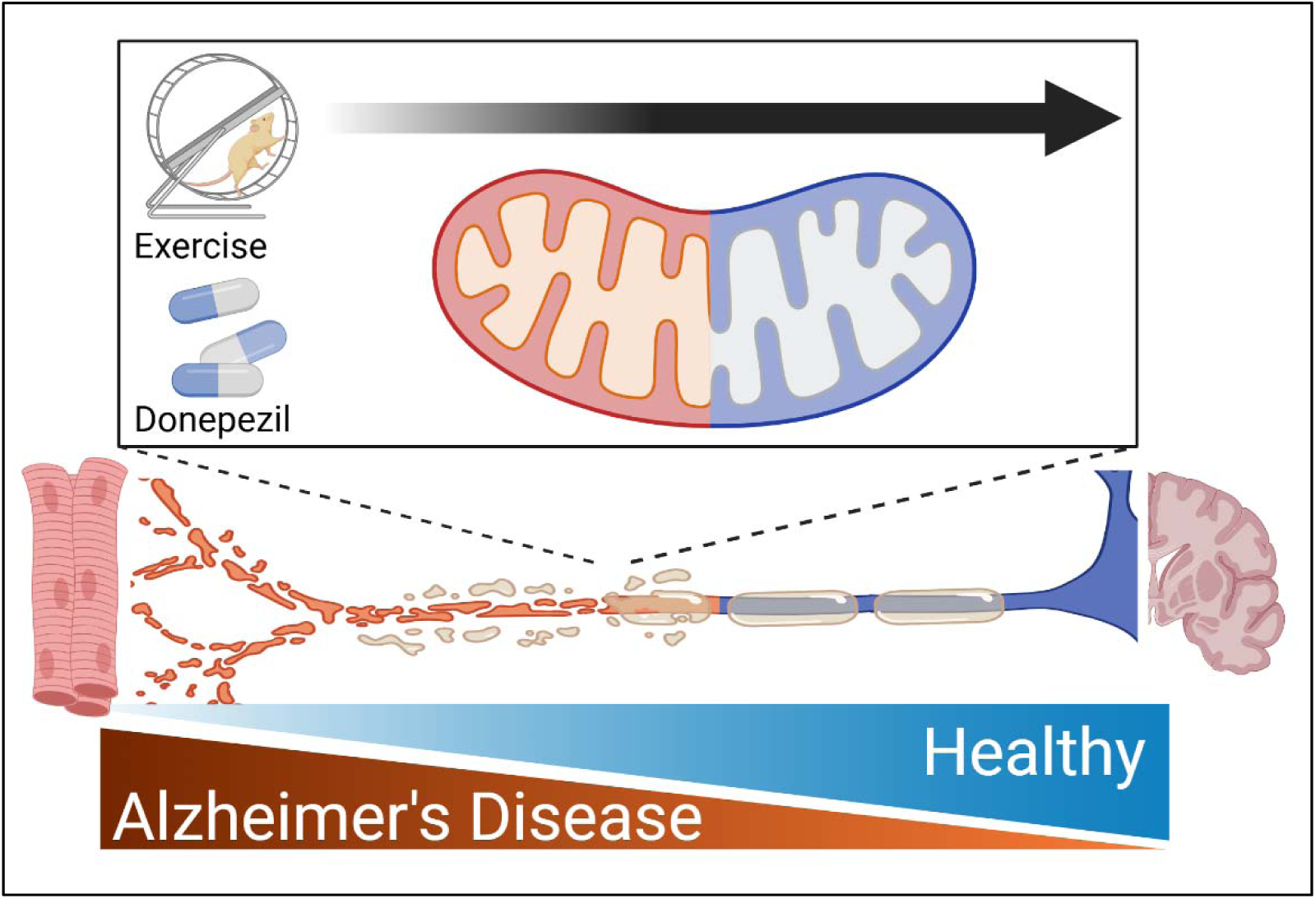

## Background

The pathogenesis of Alzheimer’s disease (AD) progresses along a continuum, with a preclinical phase that can span years to possibly decades before manifestation as mild cognitive impairment (MCI) and eventually clinical AD diagnosis^1,2^. A better understanding of phenotypes and related mechanisms of preclinical AD may bring to light novel early diagnostic strategies and interventions. Although AD is primarily an age-related brain pathology, growing evidence indicates peripheral physiological changes, such as the deterioration of skeletal muscle mass and function that occur before noticeable cognitive decline, may serve as early indicators of AD during the preclinical period. Declines in muscle mass, strength, and innervating motor nerve conduction are linked to the severity of cognitive decline and development of MCI and AD^3–5^. Alternatively, those who retain more muscle mass and strength are less likely to develop cognitive impairment or AD^5–10^. Thus, a mechanistic understanding of early neuromuscular dysfunction in AD may improve our understanding of AD progression and offer novel avenues for disease-modifying intervention.

Depolarization of α motor neurons trigger skeletal muscle contraction by transmitting action potentials along their axons to the neuromuscular junctions of innervated muscle fibers^8^. Declines in peripheral nerve conduction speed have been observed in MCI and AD subjects^3^, thus extending the AD-neurologic phenotype to the periphery. We recently showed the 5xFAD mouse model of AD develop neuromuscular dysfunction prior to the manifestation of cognitive impairment^9^. The 5xFAD mouse is a well-established model of AD that develop substantial extracellular amyloid-beta plaques by 4 months of age, with cognitive decline becoming apparent around 6 months of age ^10,11^. We found 5xFAD mice develop impaired tibial nerve-stimulated plantar flexion torque beginning at 4 months of age, as well as impaired sciatic nerve compound motor nerve action potential (CNAP)^9^. Altered nerve-to-muscle communication in early AD may partly explain the inconsistent efficacy of interventional strategies, like exercise to slow disease progression^12,13^. However, what underlies peripheral nerve dysfunction in AD or related models, like 5xFAD, is unknown.

Donepezil is a FDA approved drug used to treat cognitive impairment in AD^14–16^ that inhibits acetylcholinesterase, preventing breakdown of acetylcholine in the synaptic junction^17^. In isolated skeletal muscle, donepezil increased tetanic torque production and endplate potential duration, suggesting treatment with donepezil may have peripheral benefits on neuromuscular function during early AD^17^. This notion is supported by evidence that donepezil improves gait speed in individuals with mild AD^18^. Additionally, MCI subjects taking donepezil had normal ADP-stimulated mitochondrial respiration in skeletal muscle compared to impaired ADP-stimulated respiration in MCI subjects not on donepezil^19^.

The purpose of this study was to characterize age-related changes to the sciatic proteome in 5xFAD mice and test if exercise or donepezil could delay the initial decline in neuromuscular function seen at 4 months of age. We hypothesized either voluntary wheel running, or donepezil treatment would delay the manifestation of neuromuscular dysfunction in 5xFAD mice at 4 months of age. Understanding mechanism(s) behind neuromuscular dysfunction prior to cognitive impairment in a mouse model of AD and capacity to delay their manifestation may reveal new means to identify early signs of AD for intervention.

## Methods

### Animals

All experimental procedures were approved by the Institutional Animal Care and Use Committee (IACUC) at Virginia Tech. Male mice were housed in temperature-controlled (21°C) quarters with 12:12 hour light-dark cycle and *ad libitum* access to water and chow (Purina). Heterozygous 5xFAD mice were obtained commercially (Jackson Laboratories, strain #034840). The 5xFAD mouse overexpress five familial AD mutations [APP = Swedish (K670N, M671L), Florida (I716V), and London (V717I); PS1 = M146L and L286V) in forebrain neurons controlled by a *Thy1* promoter^11^. Male heterozygous 5xFAD mice and wild type littermates were obtained from heterozygous breeding pairs maintained in house. Genotype was confirmed via PCR.

### Voluntary Wheel Running

Heterozygous 5xFAD and WT littermates were randomly assigned to voluntary wheel running or sedentary groups at 3 months of age. Mice chosen for voluntary wheel running were individually housed in cages with running wheels. Sedentary mice were group housed in cages between 2-5 mice per cage without running wheels. Voluntary wheel running mice were given free access to a running wheel for the duration of the study^9^. All mice were provided food and water *ad libitum*.

### Exercise Capacity

After 4 weeks of access to voluntary wheel running (4 months of age) all mice (including sedentary group) were acclimated for 10 min at 13m/min at 0% incline for 3 consecutive days prior to testing endurance capacity, as previously described^9,20,21^. For voluntary running mice, running wheels were locked the night before testing. On the day of testing, mice performed an exhaustive exercise test on a treadmill set to 5% incline. Mice ran at 5 m/min for 2 min, after which treadmill speed was increased by 5 m/min every 2 min, until exhaustion. Exhaustion was defined as refusal to run, despite prodding the mice with a brush at the back of the treadmill for 20 sec.

### Donepezil Treatment

Apple-flavored donepezil and placebo treats were purchased from Bioserve. Donepezil treats were compounded with a dosage of 3mg/kg, resulting in 0.075mg of donepezil per treat, based on an average male mouse body weight of 25 grams. Mice in the donepezil or placebo groups were individually housed and given treats daily at 17:00 hrs (2 hours prior to dark cycle) for 4 weeks until sacrifice.

### In-vivo Muscle Function

Torque-frequency relationship of the plantar flexor muscles was assessed at 4 months of age, as previously described^9^. Briefly, torque via plantar flexion was measured by the torque (mN) applied to the foot pedal, and multiplied by the length of the foot pedal (m). The units were set to mN × m. Isometric torque was determined with the foot at 90° to the tibia (neutral position) over a series of stimulations at frequencies: 1, 10, 30, 50, 65, 80, 100, 120, 150, 180, and 200 Hz via F-E2 platinum-tipped needle electrodes (Natus Manufacturing, Gort, Ireland) to deliver stimulations from the 701C stimulator (ASI, Aurora, ON, Canada) via tibial nerve depolarization. To do so, two electrodes were taped together ∼3 mm apart and inserted percutaneously, distal from the knee, parallel along the tibia. Data were plotted as torque normalized to the mass of the right hind limb plantar flexors, i.e. triceps surae (mN × m/mg). Relative torque data were calculated as each respective mouse value for each time point / baseline torque × 100%.

### Sciatic Nerve Compound Motor Neuron Action Potential (CNAP)

The night prior to CNAP measures, all running wheels were locked, donepezil and placebo mice received their treat, and all mice were fasted with *ad libitum* water. Mice were anesthetized with 3% isoflurane, and a unilateral sciatic nerve was exposed for stimulation as previously described^9^. Compound nerve action potential (CNAP) duration, an index of conduction in nerve bundles^22^ was determined by a stimulus electrode placed at distance greater than 3 mm from a bipolar sensing electrode in differential amplifier mode with electrode spacing at 2 mm (2×10-3 m). The common ground for the bipolar sensing electrode and stimulation electrode was placed in the foot of the mouse. Signal from the bipolar electrode was amplified by a Hugo Sachs Electonik D-79232 amplifier and digitized by a Powerlab 4/35 at a sampling rate of 4kHz. The left sciatic nerve was stimulated at a period of 1 second, 10 mV, and 1 msec monophasic pulse using a model 4100 isolated high-powered stimulator (A-M Systems, Sequim, WA, USA).

### Blood Plasma Biomarker Analysis

Blood was collected into serum separator tubes as previously described^19^ and spun down to separate plasma and serum, which was frozen at −80°C until further analysis. Neurofilament light (NFL), were measured on a Simoa HD-X machine in plasma using the Neurology 4-Plex E kit (Quanterix)

### Tissue collection

Following Sciatic Nerve CNAP measures, blood was drawn via cardiac puncture. After which mice were sacrificed via cervical dislocation and tissues were collected and snap frozen in LN_2_.

### Western Blotting

SNAP frozen Hippocampi were cryo-homogenized via mortar and pestle and prepared for SDS PAGE as previously described^23^ membranes were probed for Amyloid beta using Biolegend Purified anti-β-Amyloid, 1-16 Antibody (# SIG-39320)

### Quantitative Proteomics of Sciatic Nerve

Sciatic nerves from both hind-limbs were collected from five control and five 5xFAD mice at 3-, 4-, and 7-months of age. For proteomics analysis, sciatic nerves were analyzed using the following protocol with minor modifications^24^. Proteins were extracted with proteomic lysis buffer [PLB1: 7 M urea, 2 M thiourea, 0.4M tris pH 8, 20% acetonitrile, 10 mM tris (2-carboxyethyl) phosphine (TCEP), 40mM chloroacetamide]. The samples were disrupted via tissue tearing and sonication. After centrifugation, the supernatants were collected to determine protein concentration via Bradford assay. 100 μg total protein were prepared from each sample and a pooled sample was created for bridging across Tanden Mass Tag (TMT) experiments. Samples were diluted five-fold with water and then trypsin digested using a 1:40 ratio of trypsin to total protein. Samples were incubated overnight for 16 hrs at 37℃. After incubation, samples were frozen at -80℃ and died in a vacuum concentrator. Each sample was cleaned with a 1 cc Waters Oasis MCX cartridge (Waters Corporation, Milford, MA), and the eluate dried *in vacuo*. Samples were resuspended in water and peptide concentration determination by BCA assay. Equal peptide amounts for each sample were further cleaned and desalted on a C18 column. The digested samples were labeled with TMT16plex Isobaric Label Reagent (Thermo Scientific) in accordance with the manufacture’s specifications. After TMT labeling, all the samples were multiplexed together into a new 1.5 mL microfuge tube. The TMT sample was dried down *in vacuo.* The samples were offline fractionated as described previously^25^. Each fraction was desalted on a C18 stage tip and reconstituted for analysis. Samples were separated on an Easy nLC-1000 UHPLC and analyzed by tandem mass spectra in a data-dependent manner with a Thermo Fisher Oritap Eclipse mass spectrometer. In brief, we processed the MS peptide spectra using Sequest (Thermo Fisher Scientific, San Jose, CA, in ProteomeDiscoverer 2.2). The mouse (taxonID 10090) Universal Proteome target protein sequence database was downloaded from UniProt (www.uniprot.org/) on August 8^th^, 2024, and was merged with a common lab contaminant protein database (http://www.thegpm.org/cRAP/index.html). The digestion enzyme was trypsin; the fragment ion mass tolerance was 0.08 Da and the precursor tolerance was 15 ppm. We set the variable modifications for oxidation of methionine (+15.9949), pyroglutamic acid conversion from glutamine (17.0265), deamidation of asparagine (+0.9840), protein N-terminal acetylation (+42.0106) and TMT 16plex (+229.1629) modification of lysine and peptide N-terminus. We specified carbamidomethyl of cysteine as a fixed modification. We used the Percolator algorithm with a concatenated target-decoy database approach to control the false discovery rate (https://doi.org/10.1038/nmeth1113). We reported protein lists with a 1% FDR threshold. Reporter ion intensities were adjusted by correction factors in all samples according to the algorithm described in i-Tracker78 according to the TMTpro 16plex Lot Number WF324548 product data sheet from ThermoFisher Scientific. Normalization was performed iteratively (across samples and spectra) on intensities, as described in statistical analysis of Relative Labeled Mass Spectrometry Data from Complex Samples Using ANOVA.79 Pooled protein samples from all samples in the two TMT experiments were used for normalization, using two TMT channels for each experiment. Medians were used for averaging. Spectra data were log-transformed, pruned of those matched to multiple proteins, and weighted by an adaptive intensity weighting algorithm. Differentially expressed proteins were determined by applying a permutation test with an unadjusted significance level p < 0.05 corrected by Benjamini-Hochberg. We set the hypothesis test method to t-test, and we reported p-values that were adjusted using the Benjamini-Hochberg correction for false discovery rate.

### Quantitative Proteomics of Sciatic Nerve from exercise and donepezil treated mice

Exercised and donepezil treated mice were generated as described above. Similar to the quantitative proteomics of Sciatic Nerve data described above, sciatic nerve samples from 4-month-old exercise and donepezil treated mice were collected. Proteins were extracted with proteomic lysis buffer [ PLB1: 7 M urea, 2 M thiourea, 0.4M tris pH 8, 20% acetonitrile, 10 mM tris (2-carboxyethyl) phosphine (TCEP), 40mM chloroacetamide]. The samples were disrupted via tissue tearing and sonication. After centrifugation, the supernatants were collected to determine protein concentration via Bradford assay. 100 μg was prepared from each sample and a pooled sample was created for bridging across TMT experiments. Samples were diluted five-fold with water and then trypsin digested using a 1:40 ratio of trypsin to total protein. Samples were incubated overnight for 16hrs at 37C. After incubation, samples were frozen at -80C and died in a vacuum concentrator. Each sample was cleaned with a 1 cc Waters Oasis MCX cartridge (Waters Corporation, Milford, MA), and the eluate dried *in vacuo*. Samples were resuspended in water and peptide concentration determination by BCA assay. Equal peptide amounts for each sample were desalted on a C18 column. The digested samples were labeled with TMT16plex Isobaric Label Reagent (Thermo Scientific) in accordance with the manufacture’s specifications. After TMT labeling, all the samples were multiplexed together into a new 1.5 mL microfuge tube. The TMT sample was dried down *in vacuo.* The samples were offline fractionated as described previously^25^. Each fraction was desalted on a C18 stage tip and reconstituted for analysis. Data were collected as described above.

### Heatmap and pathway analysis of proteomics data

Pathway analysis, hierarchical clustering, and heat map production were carried out as previously described ^11–17^. The hierarchical clustering and heat maps were generated using Cluster 2.0 from the Eisen Laboratory modified by Michiel de Hoon (http://bonsai.hgc.jp/∼mdehoon/software/cluster/). Java Tree Viewer was used to view and color the heat map. Identification of KEGG and GO pathways were determined via ShinyGO v0.82 (https://bioinformatics.sdstate.edu/go/) and cross referenced using PANTHER (https://pantherdb.org/).

### Statistics

Data are presented as mean ± SEM and were analyzed and plotted via GraphPad Prism 9.5.1. Data were analyzed via Student’s t-test when one variable was present, two-way ANOVA when two variables were present, or repeated measures two-way ANOVA when two variables were measured in the same animals over time. In groups with significant standard deviation differences, data were transformed to further detect significance among groups. Post-hoc analyses were only performed when a significant interaction between a categorical and a quantitative variable was found, which are indicated in the figures where applicable. Statistical significance was established *a priori* as p < 0.05.

## Results

### 3.1 Cross Sectional Age Time Course Assessment of Sciatic Nerve Proteome

We previously demonstrated tibial nerve-stimulated hindlimb skeletal muscle torque production declines at 4 months of age in 5xFAD mice due to impaired peripheral nerve function^9^, analogous to findings in MCI and AD subjects^3^. Impairments in nerve transmission within the brain are known in 5xFAD mice^26–28^ and in humans with AD^29–31^. However, to our knowledge, no analysis of the peripheral nerve proteome along the functional spectrum in an AD-like model has been performed. To assess age-related proteome changes in sciatic nerve of 5xFAD mice, we performed TMT-labeled proteomic analysis on sciatic axon sections from male 5xFAD mice and WT littermates at 3, 4, and 7 months of age (Fig. 1A). Ages were chosen to represent when nerve-to-muscle communication is physiologically equivalent between groups (3 months), when neuromuscular dysfunction first appears (4 months), and at an advanced stage (7 months)^9^. At 3 months, we observed 223 significantly abundant proteins upregulated in WT sciatic axons compared to 72 in 5xFAD (Fig. 1B). KEGG Pathway analysis of significantly enriched proteins revealed they were related to contraction and calcium signaling pathways (Fig. 1B), although we previously observed equivalent nerve-stimulated skeletal muscle torque production and sciatic CNAP between 5xFAD and WT at 3 months of age^9^. At 4 months of age, corresponding to the initial decline in nerve-to-muscle communication^9^, the 5xFAD sciatic proteome underwent a pronounced remodeling with 329 significantly abundant proteins enriched in the 5xFAD compared to 159 in WT (Fig. 1C). KEGG Pathway analysis of significantly enriched proteins indicated nitrogen and amino acid metabolism were altered (Fig. 1C), potentially alluding to changes in neurotransmitter regulation. At 7 months, the sciatic proteome was again skewed towards WT with 143 abundant proteins enriched compared to 54 in 5xFAD (Fig. 1D). KEGG Pathway analysis of significantly enriched proteins revealed they were related to fatty acid and amino acid metabolism (Fig. 1D). As the remodeling of the sciatic proteome in 5xFAD mice mirrored the previously observed decline in nerve-stimulated muscle torque production between 3 and 4 months of age^9^, we compared the 3-and 4-month sciatic proteomes in 5xFAD mice. The 5xFAD sciatic proteome at 4 months of age had 210 abundant proteins compared to 114 at 3 months of age that related to amino acid and cholesterol metabolism, as well as mitophagy (Fig. 1E), suggesting mitochondrial dysfunction is a key event coincided with neuromuscular dysfunction at 4 months of age. To determine how specific sets of proteins in sciatic axons were differentially enriched with age in 5xFAD mice compared to same aged WT littermates, enriched proteins were organized by hierarchical clustering and K-means statistics (Fig. 1F). Remodeling of the sciatic proteome was most apparent in 5xFAD mice at 4 months of age with enrichments in proteins related to Schwann cell remodeling, calcium channel activity, axon remodeling, lipid/glucose metabolism, synaptic vesicle fusion, mitochondria remodeling, and amino acid metabolism (Fig. 1F; middle row). Together, these data suggest that onset of neuromuscular impairment at 4 months of age in 5xFAD mice^9^ corresponds to sudden remodeling in the proteome of the sciatic nerve, particularly in pathways that converge on mitochondria. Thus, the 3–4-month age period in 5xFAD mice may represent an interventional window that has translational relevance as peripheral nerve dysfunction has been noted in MCI and AD subjects^3^.

**Figure 1.**
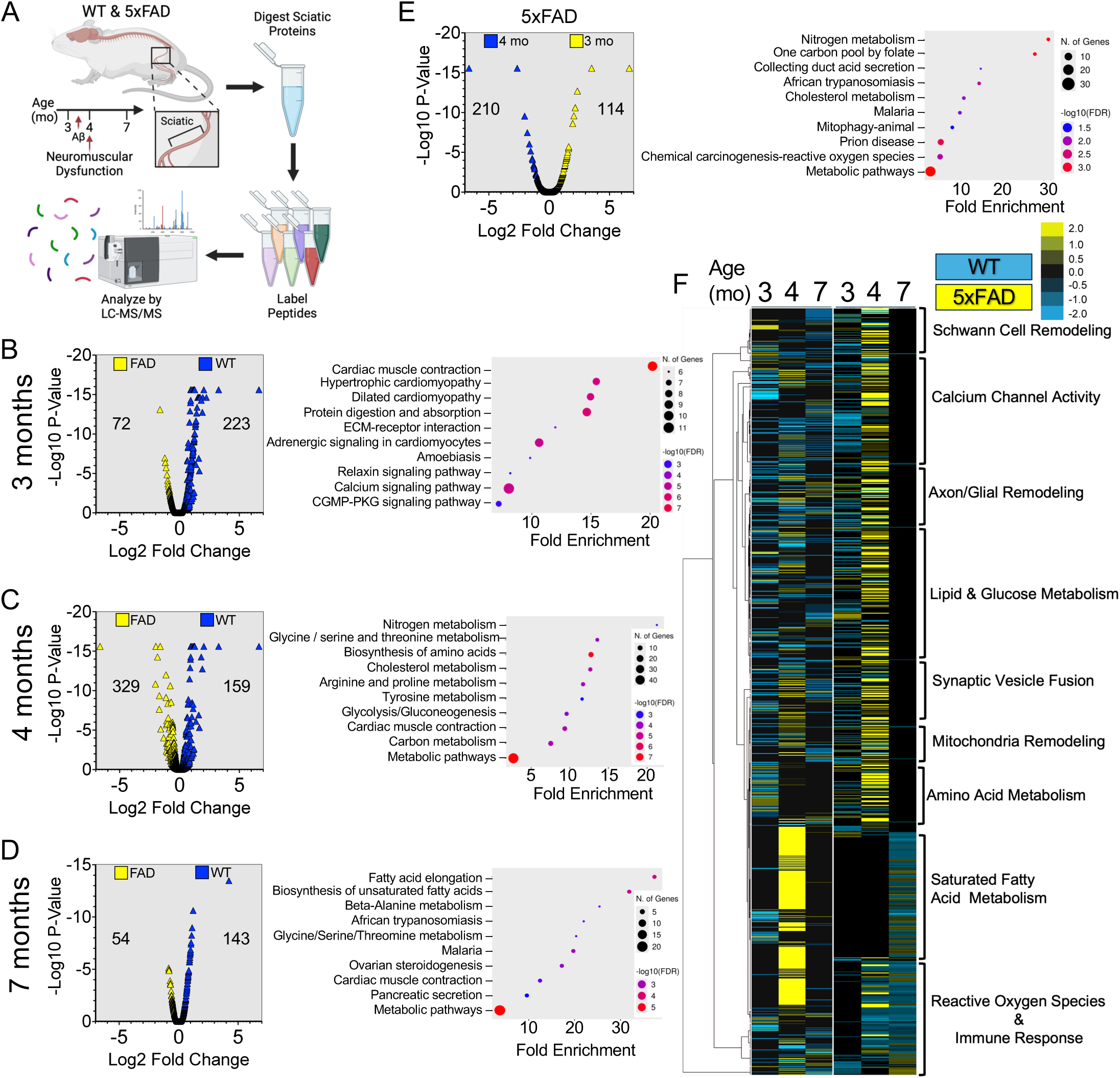
Cross sectional age time course assessment of sciatic nerve proteome in 5xFAD mice: (A) Schematic outlining proteomic profiling of sciatic nerve from 5xFAD mice at 3, 4, and 7-months of age. (B) Volcano plot of 3-month-old WT and 5xFAD sciatic with corresponding KEGG pathway analysis of statistically significant proteins. (C) Volcano plot of 4-month-old WT and 5xFAD sciatic with corresponding KEGG pathway analysis of statistically significant proteins. (D) Volcano plot of 7-month-old WT and 5xFAD sciatic with corresponding KEGG pathway analysis of statistically significant proteins. (E) Volcano plot of 3-4-month-old 5xFAD sciatic with corresponding KEGG pathway analysis of statistically significant proteins. (F) Heatmap and K-means statistical analysis of proteoms of WT and 5xFAD sciatic nerves. Left: hierarchical clustering of proteins significantly different between 3, 4, and 7-month fractions (threshold for clustered proteins was determined by significance between groups). Right: proteins were further grouped using k-means statistics, breaking the 3, 4, 7-month significant genes into 10 protein clusters. n = 5 per age and genotype.

### 3.2 Voluntary Wheel Running Attenuates Neuromuscular Impairment at 4 Months of Age in 5xFAD Mice

We have shown exercise training that went to 6 months of age in 5xFAD mice resulted in an altered adaptive response in skeletal muscle^9^, suggesting a threshold where the pathological burden cannot be overcome by exercise. We hypothesized exercise begun prior to neuromuscular dysfunction may be able to delay onset of nerve-stimulated skeletal muscle torque loss at 4 months of age in 5xFAD mice. 5xFAD mice and their WT littermates were given voluntary wheel running access for 4 weeks starting at 3 months of age (Fig. 2A). To assess whether voluntary wheel running began at 3 months of age could prevent declines in skeletal muscle torque production we assessed torque production of the plantar flexors (i.e. gastrocnemius, plantaris, and soleus) via stimulation of the tibial nerve *in-vivo*. Voluntary wheel running significantly improved plantar flexion torque production in WT, as evidenced by increase in the force frequency curve normalized to the wet weight of the triceps surae (Fig. 2B). At 4 months of age, sedentary 5xFAD had a lower force frequency profile compared to WT, as we had observed previously^9^ (Fig. 2B, C). Interestingly, exercise training maintained 5xFAD nerve-stimulated torque frequency responses comparable to WT sedentary animals but there was no improvement beyond sedentary WT as seen in WT exercised mice (Fig. 2B, C). Three days after testing nerve-stimulated skeletal muscle torque, exercise capacity was tested (Fig. 2A). Exercise capacity significantly increased with voluntary wheel running in both WT and 5xFAD mice (Fig. 2,B), as reported previously^9^. This outcome suggests that the difference in nerve-stimulated skeletal muscle torque frequency between WT and 5xFAD mice is not due to a difference in exercise performance capacity. To further examine the effect of voluntary wheel running on neuromuscular function, we assessed sciatic CNAP *in vivo* three days following exercise capacity testing (Fig. 2A). There was no effect of voluntary wheel running seen in WT Ex mice (Fig. 2E), suggesting a possible functional ceiling in CNAP in the motor neuron with beneficial effects seen in torque production resulting from adaptation at the neuromuscular junction^32^. Sedentary 5xFAD mice had significantly slower sciatic CNAP compared to WT (Fig. 2E), analogous to our previous findings^9^. However, exercise training mitigated the increase in sciatic CNAP in 5xFAD at 4 months of age (Fig. 2E). Taken together these findings demonstrate exercise can attenuate the initial decline in neuromuscular function at 4 months of age in 5xFAD mice.

**Figure 2.**
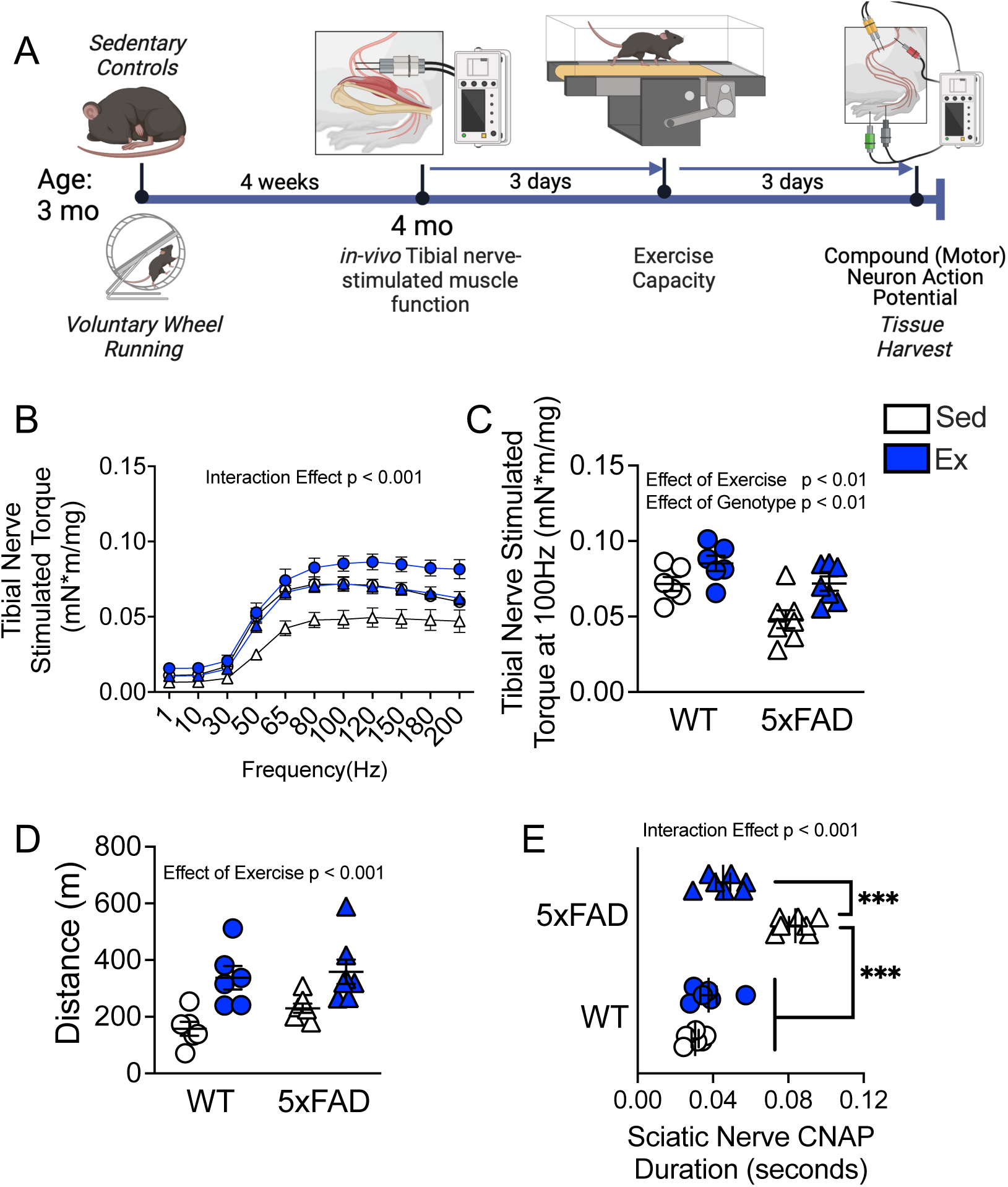
Voluntary wheel running attenuates neuromuscular impairment at 4 months of age in 5xFAD mice: (A) Schematic outlining study design of voluntary wheel running intervention. (B) Tibial nerve-stimulated torque frequency of plantar flexors. (C) Tibial nerve stimulated torque at 100Hz. (D) Exercise capacity. (E) In vivo compound nerve action potential (CNAP) of sciatic nerve. n= 6 for all groups. Data presented as mean ± SEM and repeated measures two-way ANOVA was performed and Tukey’s post-hoc test performed when significant interaction between variables was observed (∗ = P < 0.05, ∗∗ = P < 0.01, and *** = P<0.001). Panel A created in Biorender.

### 3.3 Proteomic Profiling of The Sciatic Nerve Following Exercise Training

To determine how exercise attenuated neuromuscular decline in 4-month-old 5xFAD mice, we performed TMT-labeled proteomics of sciatic nerve axon sections. Following LC-MS/MS analysis of sciatic nerves, we identified 7,982 proteins from a total of 18,636 unique peptide spectra at a 2-peptide and a 1.1% FDR minimum. Of these 7,982 proteins, 33 were significantly enriched in 5xFAD-Sed compared to 216 in WT-Sed, related to propanoate and amino acid metabolism, as well as PPAR signaling (Fig. 3A). this data indicates that the initial metabolic insult is present prior to the decline observed at 4 months of age. In 5xFAD exercised sciatic, 144 proteins were enriched compared to 83 in sedentary counterparts, related primarily to calcium and contraction signaling (Fig. 3B). Because exercise training-maintained nerve-stimulated torque frequency and sciatic CNAP comparable to WT sedentary animals (WT-Sed), we compared 5xFAD exercise sciatic proteome to WT-Sed. In 5xFAD exercised mice, 88 proteins were enriched in sciatic compared to 143 in WT-Sed related to fatty acid metabolism and calcium signaling (Fig. 3C). While the comparison between 5xFAD and WT-Sed do not overlap 1:1, this data indicated that exercise stabilized mitochondrial and metabolic proteins dysregulated by the disease etiology of 5xFAD mice at 4 months of age. Further analysis via hierarchical and K-means clustering showed distinct differences between voluntary wheel running and genotype (Fig. 3D, Fig. S1). As exercise training maintained neuromuscular function comparable to WT sedentary mice (Fig. 2), we focused on proteins whose expression was different between WT and 5xFAD sedentary animals, inversely affected by exercise in 5xFAD mice compared to sedentary, and then roughly equivalent with WT sedentary following exercise training in 5xFAD. A protein-protein interaction string map was generated from identified proteins (Fig. S2) and interacting clusters were identified (Fig. 3E, S2). Identified protein-protein interaction clusters were related to inflammation (Apcs, Hp, Saa1, Saa2, Saa4, Crp), calcium buffering (Asph, Jsrp1, Myoz1, Tcap, Srl, Hrc), and mitochondrial turnover (Orm2) (Fig. 3E, F). These data suggests that exercise, when begun prior to neuromuscular dysfunction, attenuates neuromuscular dysfunction in 5xFAD mice by suppressing an inflammatory cascade that coincides with increased mitochondrial turnover at 4 months of age, potentially maintaining calcium buffering capacity (Fig. 3F).

**Figure 3.**
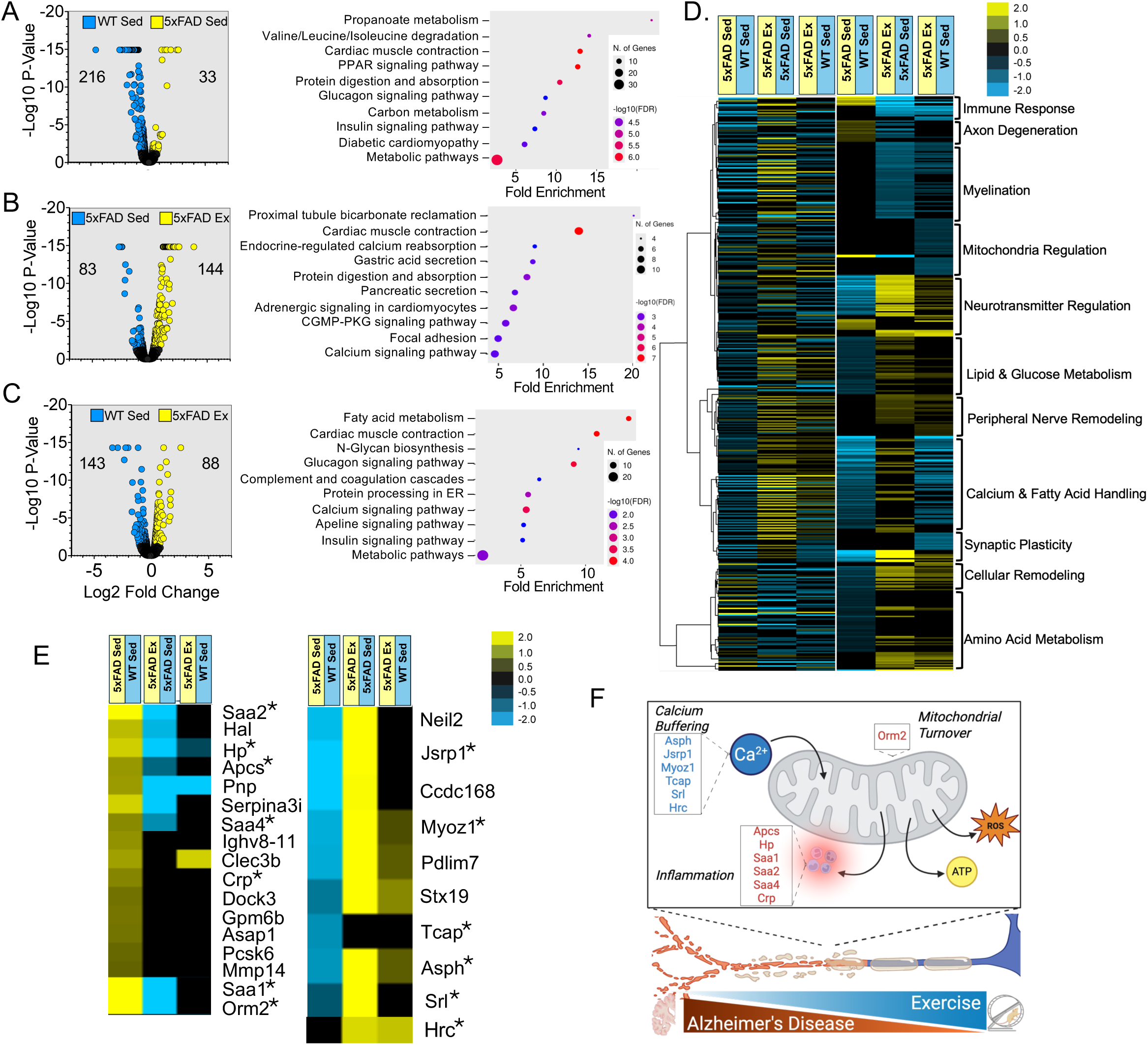
Proteomic profiling of sciatic nerve following exercise training: (A) Volcano plot of WT Sed and 5xFAD Sed sciatic and KEGG annotation. (B) Volcano plot of 5xFAD Sed and 5xFAD Ex sciatic with corresponding KEGG pathway analysis of statistically significant proteins. (C) Volcano plot of WT Sed and 5xFAD Ex sciatic with corresponding KEGG pathway analysis of statistically significant proteins. (D) Heatmap and K-means statistical analysis of proteomes from WT and 5xFAD sciatic nerves. Left: hierarchical clustering of proteins significantly different between WT and 5xFAD Sed and Ex (threshold for clustered proteins was determined by significance between groups). Right: proteins were further grouped using k-means statistics, breaking the significant genes into 10 protein clusters. (E) Heat map of proteins inversely affected by exercise and returned to WT levels. * denotes proteins that group together in STRING analysis. (F) Schematic of protein-protein interacting clusters, representing processes maintained to WT Sed by exercise in 5xFAD.

We next assessed if maintained sciatic proteome and neuromuscular function in 5xFAD mice following exercise training at 4 months of age coincided with improved indices of AD-like neuropathology. Circulating neurofilament light chain (NfL) in plasma was significantly elevated in both sedentary and exercised 5xFAD mice compared to WT counterparts (Fig. S3A). Expression of high molecular weight Aβ oligomers in hippocampus lysates were also elevated in both sedentary and exercise trained 5xFAD mice (Fig. S3B). Together, these data suggest that attenuation of neuromuscular dysfunction in 5xFAD mice can result from exercise training independent of improvement in markers associated with the AD-like phenotype of the model.

### 3.4 Donepezil Attenuates Neuromuscular Impairment at 4 Months of Age in 5xFAD Mice

As the efficacy of exercise as an interventional strategy in AD is dependent on a variety of factors, such as timing along the continuum of the pathology^9,12,33^, we tested whether a currently approved therapeutic may also delay neuromuscular decline in 5xFAD mice. The acetylcholinesterase inhibitor Donepezil is FDA approved to slow cognitive impairment in AD^14,15^. Donepezil has been shown to maintain mitochondrial respiration in skeletal muscle in humans with MCI^19^ as well as gait speed^18^. As sciatic proteome changes at 4 months of age in 5xFAD mice involves mitochondria-centric processes (e.g. mitochondrial turnover, calcium buffering, and metabolism) (Fig. 1F, 3D-F) , we hypothesized treatment with donepezil could delay onset of neuromuscular dysfunction. Following 4 weeks of daily donepezil treatment, beginning at 3 months of age, we assessed *in-vivo* tibial nerve-stimulated plantar flexor torque production (Fig. 4A). Donepezil prevented decline of plantar flexion torque production in 4-month-old 5xFAD mice compared to placebo (PLCB) treated mice when normalized to the wet weight of the triceps surae (Fig. 4B, C). Decline in sciatic nerve CNAP *in-vivo* was attenuated in 5xFAD mice treated with donepezil compared to placebo but was still significantly slower compared to WT sedentary animals (Fig. 4C). In sum, donepezil treatment begun before neuromuscular dysfunction in 5xFAD mice can slow its development.

**Figure 4.**
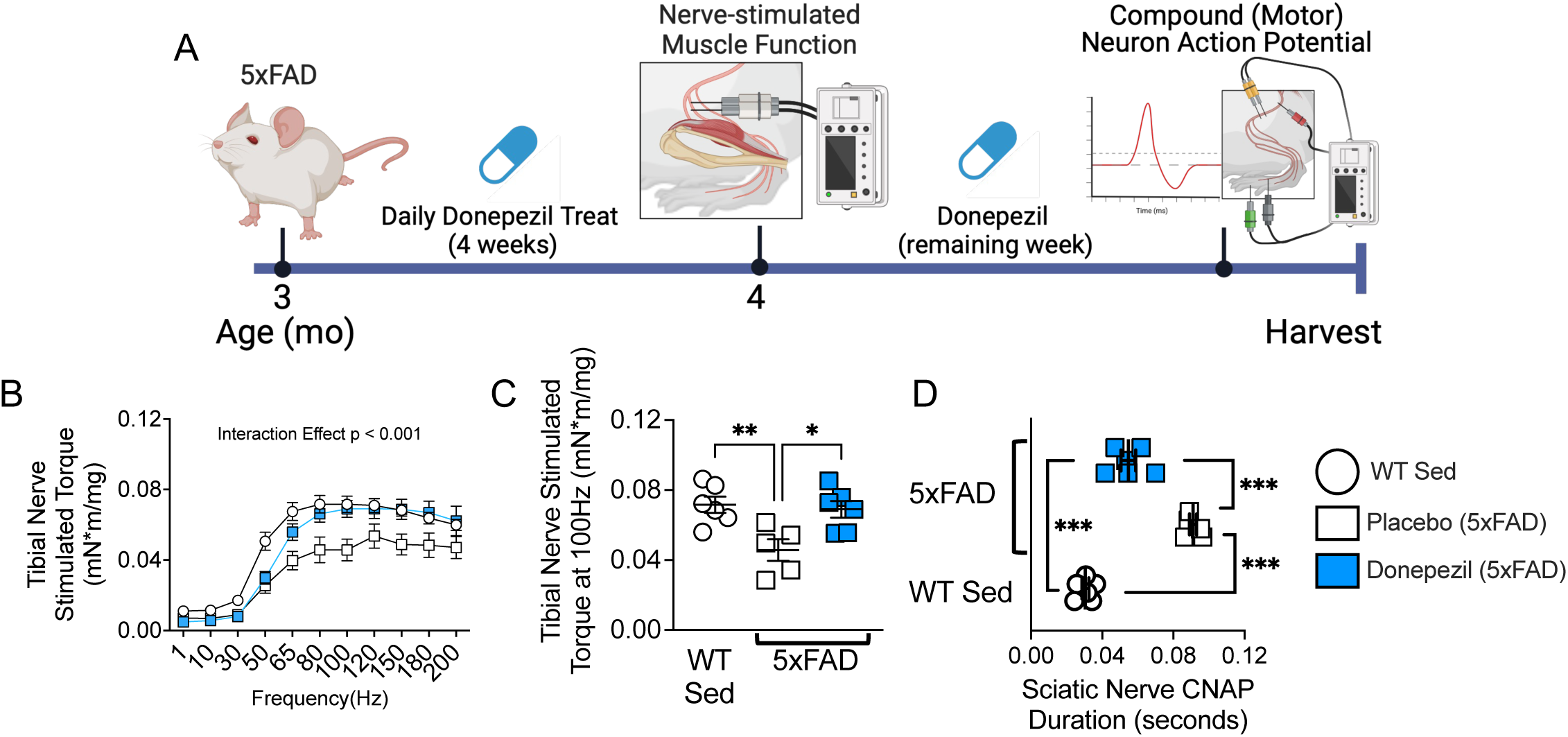
Donepezil attenuates neuromuscular impairment at 4 months of age in 5xFAD mice: (A) Study design of donepezil treatment. (B) Tibial nerve-stimulated skeletal muscle torque frequency. (C) Tibial nerve stimulated torque at 100Hz. (D) In vivo compound nerve action potential (CNAP) of sciatic. n= 6 for WT, n=6 for donepezil and n=5 for placebo. Data presented as mean ± SEM and repeated measures two-way ANOVA was performed and Tukey’s post-hoc test performed when significant interaction between variables was observed (∗ = P < 0.05, ∗∗ = P < 0.01, and *** = P<0.001). Panel A created in Biorender.

### 3.5 Proteomic Profiling of The Sciatic Nerve in Response to Donepezil Treatment

To determine how donepezil treatment attenuated neuromuscular dysfunction in 4-month-old 5xFAD mice, we again performed TMT-labeled proteomics of sciatic nerve axon sections. Following LC-MS/MS analysis of sciatic nerves from donepezil treatment, we identified 5,861 proteins from a total of 61,925 unique peptide spectra at a 2-peptide and a 0.5% FDR minimum. Of these 5,861 proteins, 164 were significantly enriched in 5xFAD-PLCB compared to 54 in WT-Sed (Fig. 5A), related to nitrogen metabolism, amino acid metabolism, contraction, and PPAR signaling pathways, which overlaps with equivalent comparisons between 4-month-old 5xFAD and WT (Fig. 1A) as well as 5xFAD sedentary mice (Fig. 3C). In 5xFAD treated with donepezil, 91 proteins were enriched compared to 84 in placebo treated, related to nitrogen metabolism, protein metabolism, contraction and PPAR signaling pathways (Fig. 5B). In 5xFAD treated with donepezil, 33 proteins were enriched compared to 193 in WT-Sed, related to fatty acid metabolism, complement and coagulation cascades (inflammation), and metabolic pathways (Fig. 5C), which were also observed in exercise trained 5xFAD mice (Fig. 3C). Heatmaps of hierarchical and K-means clustering showed distinct differences between 5xFAD-DPNZ and PLCB in relation to WT-Sed (Fig. 5D). As donepezil treatment maintained neuromuscular function similar to WT-Sed mice (Fig. 4), we again focused on proteins whose expression was different between WT and 5xFAD placebo treated animals, inversely affected by donepezil in 5xFAD mice compared to placebo, and then largely equivalent with WT sedentary following donepezil treatment in 5xFAD. Protein-protein interaction string map was generated from identified proteins (Fig. S5) and interacting clusters were identified (Fig. 5E, S4). Identified protein-protein interaction clusters were related to mitochondrial immune response (Ifit3, Bst2), inflammation (Apcs, Cfn, Saa1, Saa2, Apoc1, Hpx), and calcium buffering (Cacnb1, Hrc, Myoz3, Ryr1, Trdn, Jph1, Jph2, Trim72, Tcap, Tnnc2, Mylpf, Cavin4, Asph, Hhatl). These data suggest that donepezil, when begun prior to neuromuscular dysfunction, attenuates neuromuscular decline in 5xFAD mice by suppressing a mitochondrial immune response and inflammatory cascade that coincides with maintaining calcium buffering capacity (Fig. 5F).

**Figure 5.**
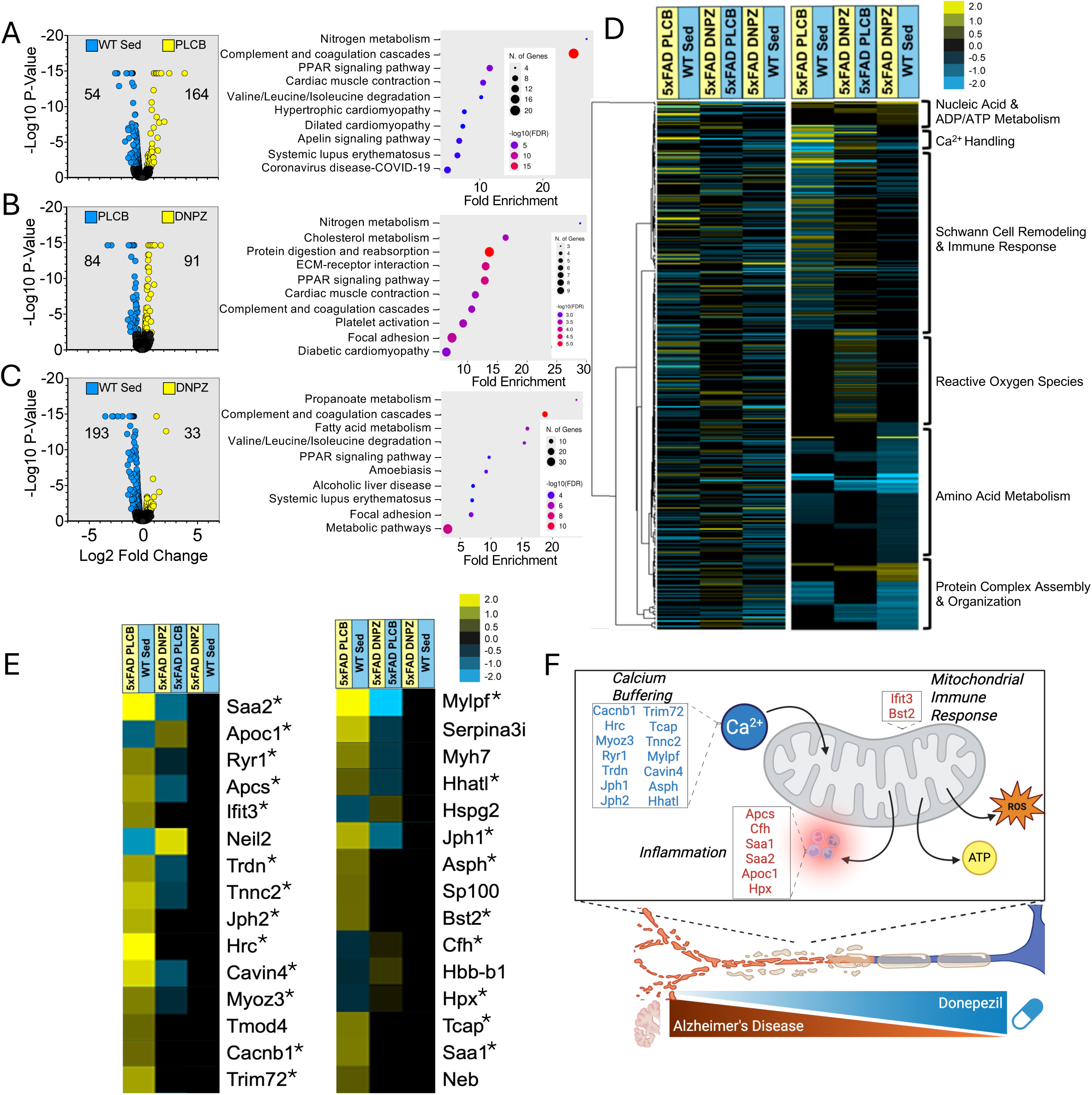
Proteomic profiling of sciatic nerve following donepezil treatment: (A) Volcano plot of WT Sed and 5xFAD placebo (PLCB) sciatic with corresponding KEGG pathway analysis of statistically significant proteins. (B) Volcano plot of 5xFAD PLCB and 5xFAD donepezil (DNZP) sciatic with corresponding KEGG pathway analysis of statistically significant proteins. (C) Volcano plot of WT Sed and 5xFAD DNZP sciatic with corresponding KEGG pathway analysis of statistically significant proteins. (D) Heatmap and K-means statistical analysis of proteomes from WT and 5xFAD sciatic nerves. Left: hierarchical clustering of proteins significantly different between WT and 5xFAD PLCB and donepezil fractions (threshold for clustered proteins was determined by significance between groups). Right: proteins were further grouped using k-means statistics, breaking the significant genes into 10 protein clusters. (E) Heat map of proteins inversely affected by donepezil and returned to WT levels. * denotes proteins that group together in STRING clusters. (F) Schematic of protein-protein interacting clusters, representing processes maintained to WT Sed by donepezil in 5xFAD.

We also assessed if maintained sciatic proteome and neuromuscular function in 5xFAD mice treated with donepezil coincided with improvements in indices of AD-like neuropathology. Plasma NfL was significantly elevated in both placebo and donepezil treated groups compared to WT sedentary (Fig. S5A). Expression of high molecular weight Aβ oligomers in hippocampus lysates were also elevated in both placebo and donepezil treated 5xFAD mice (Fig. S5B). Together, these data suggest attenuated neuromuscular decline in 5xFAD mice can be obtained with donepezil treatment independent of improvement in markers associated with the AD-like phenotype of the model.

### 3.6 Donepezil Fails to Rescue Neuromuscular Dysfunction in 5-Month-Old 5xFAD Mice

We next tested whether donepezil treatment could rescue neuromuscular function once manifested. We treated 5-month-old 5xFAD mice, an age at which neuromuscular dysfunction is already present^9^, with donepezil or placebo for 4 weeks (Fig. S6A). Following 4 weeks of treatment, at 6 months of age, we assessed tibial nerve-stimulated plantar flexor torque production *in-vivo* and direct muscle stimulation (Fig. S6A). As we observed previously^9^, nerve-stimulated skeletal muscle torque was significantly lower compared to direct muscle-stimulated torque production, again reiterating that the source of neuromuscular dysfunction was localized to the peripheral motor nerve and not to the skeletal muscle’s capacity to produce torque during contraction (Fig. S6B)^9^. Compared to placebo treated animals, donepezil did not improve tibial nerve-stimulated skeletal muscle torque production at 6 months of age (Fig. S6C). In fact, at 6 months of age, donepezil treated animals had lower direct muscle-stimulated torque production compared to placebo (Fig. S6D), suggesting donepezil may have AD-like disease stage-dependent negative effects on neuromuscular function. Furthermore, *in-vivo* sciatic nerve CNAP was not improved in 6-month-old 5xFAD mice treated with donepezil compared to placebo (Fig. S6E). While donepezil treatment attenuates the development of neuromuscular dysfunction in 5xFAD mice (Figs. 4 and 5), treatment with the same dose cannot rescue neuromuscular function once manifested.

### 3.6 Shared and Unique Proteomic Profiles of The Sciatic Nerve in Voluntary Wheel Running and Donepezil

Both exercise training and donepezil treatment in 5xFAD mice resulted in similar physiological maintenance of neuromuscular function as WT-Sed mice when begun at 3 months of age (Fig. 6A) and slowed increased CNAP (Fig. 6B). Therefore, we sought to identify both shared and unique pathways through which exercise and donepezil are protective. For this comparison we analyzed all sciatic proteins in 5xFAD exercise and donepezil treated groups that were not significantly different from WT-Sed sciatic, which resulted in 173 proteins that were attenuated to WT sedentary levels. We identified 114 proteins in common between 5xFAD exercise and donepezil groups (Fig. 6C). 5xFAD exercise mice had 38 unique proteins that were not significantly different from WT-Sed, whereas 5xFAD mice had 21 unique proteins (Fig. 6C). A protein-protein interaction string map was generated for proteins unique to exercise or donepezil to identify interacting clusters (Fig. S7). Identified protein-protein interaction clusters in 5xFAD exercise that were uniquely maintained relative to WT Sed levels were related to TCA cycle (Ampd1, Suclg2, Mccc1-2, Ivd, Dbt, Pcx) and ATP cycling (Ak2) (Fig. S8A, B). Identified protein-protein interaction clusters in 5xFAD mice treated with donepezil that were uniquely maintained relative to WT Sed levels were related to calcium handling (Asph, Ryr1, Myoz1, Micu2) and glycogen metabolism (Phkb, Phkg1, Pygm) (Fig. S8A, B). Heatmaps of hierarchical and K-means clustering showed biological processes shared between exercise and donepezil to attenuate neuromuscular dysfunction in 4-month-old 5xFAD mice (Fig. 6D). The 114 shared proteins between exercise and donepezil were involved in pathways related to trafficking and metabolism of fatty acids, synaptic transmission, and immune response (Fig. 6D, E). Thus, exercise and donepezil act, in part, by attenuating proteome changes related to mitochondrial-centric fatty acid metabolism and suggests age-dependent changes in sciatic proteome related to mitochondrial-dependent axon function underlie neuromuscular dysfunction at 4 months of age in 5xFAD mice. In sum, these findings highlight distinct and overlapping mechanisms through which voluntary exercise and a clinically utilized pharmacological intervention, donepezil, influence early sciatic proteome remodeling in 5xFAD mice to maintain neuromuscular function, thus providing insight into potential therapeutic targets and timing strategies.

**Figure 6.**
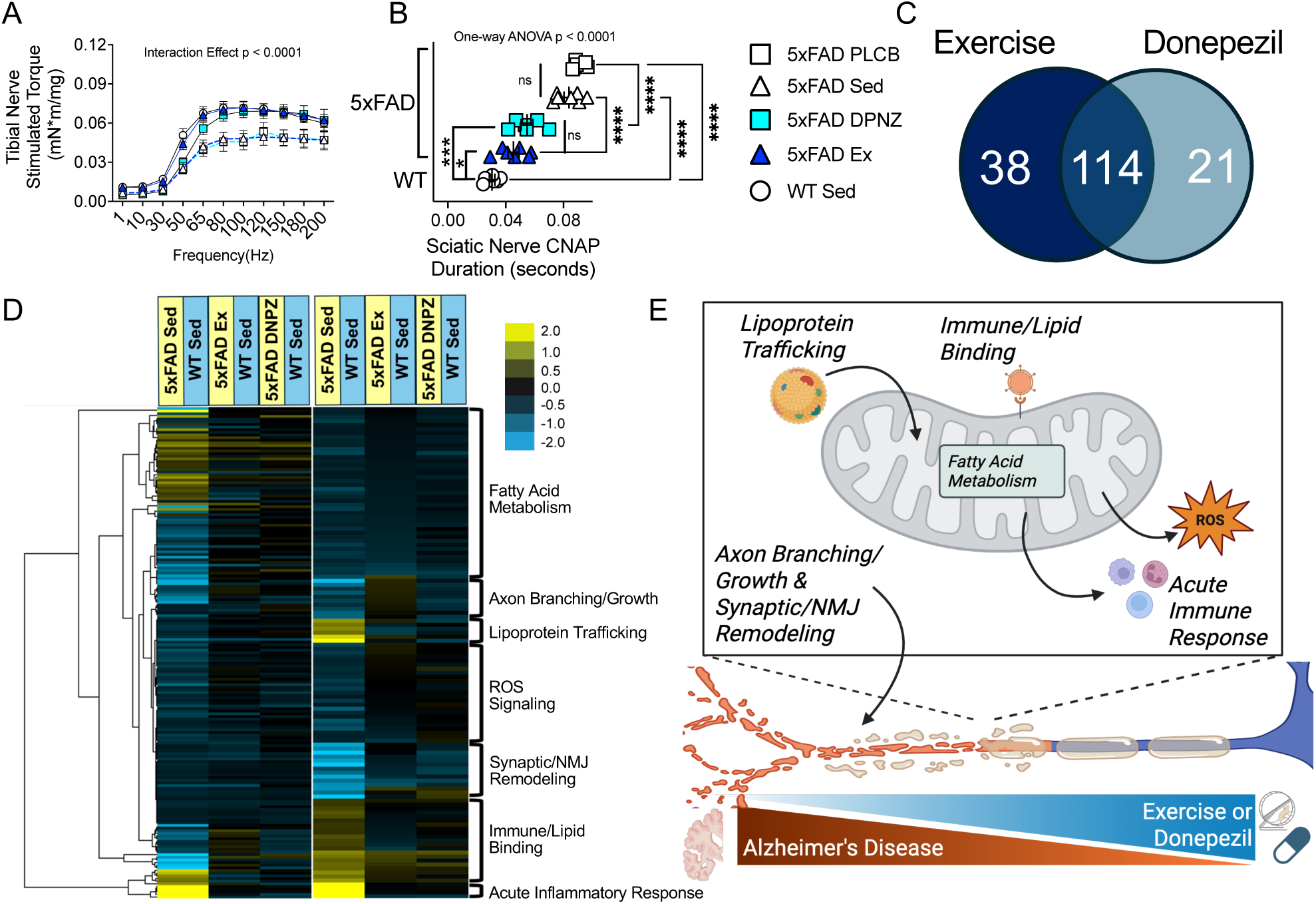
Shared proteomic changes in the sciatic nerve between exercise training and donepezil treatment: (A) Tibial nerve-stimulated skeletal muscle function in donepezil (DNZP) and voluntary wheel running (Ex). (B) CNAP in donepezil and voluntary wheel running. (C) Venn diagram of number of shared and unique proteins with voluntary wheel running and DPNZ treatment. (D) Heatmap and K-means statistical analysis of proteomes from WT and 5xFAD sciatic nerves. Left: hierarchical clustering of proteins significantly different between WT and 5xFAD PLCB and DPNZ fractions (threshold for clustered proteins was determined by significance between groups). Right: proteins were further grouped using k-means statistics, breaking the significant genes into 10 protein clusters. (E) Schematic of shared processes maintained to WT Sed levels by both Ex and DPNZ in 5xFAD.

## Discussion

In this study we investigated the proteomic profiles of sciatic nerves from 5xFAD mice, a model of AD, at 3, 4, and 7 months of age. Additionally, we investigated the effect of either voluntary wheel running or donepezil treatment initiated at 3 months of age to mitigate declines in neuromuscular function that manifest at 4 months of age. Previously, in 5xFAD mice, we discovered a decline in neuromuscular function that manifested by 4 months of age^9^. Notably, impaired neuromuscular function was due to impaired neural conduction to skeletal muscle, as evidenced by impaired nerve-stimulated muscle torque production and sciatic CNAP^9^. Our age-dependent proteomic analysis of the sciatic nerve sought to elucidate potential molecular mechanisms underlying the early neuro-muscular-degenerative processes in peripheral motor nerves. We found distinct changes in the sciatic proteome with significantly abundant proteins in pathways involving mitochondrial quality control, metabolism, calcium regulation, and synaptic transmission elevated at 4 months of age. By 7 months of age many of the proteins seen at 4 months of age were absent, suggesting an early remodeling in the sciatic proteome may drive the neuromuscular dysfunction we previously characterized^9^. Neuromuscular impairments in preclinical AD are linked with severity of cognitive decline^4–7,34,35^, but little has been done to investigate the potential mechanistic cause(s). Impairment occurs before noticeable cognitive deficits, suggesting neuromuscular dysfunction is an early characteristic of AD-like pathology, which may help explain motor deficits observed in preclinical AD^36^.

Based on the 4 month age of onset of neuromuscular dysfunction in 5xFAD mice^9^ and the associated sciatic proteome remodeling we defined herein, we hypothesized that interventions such as voluntary wheel running or donepezil initiated before the age of decline could delay neuromuscular impairment onset. Our findings revealed that voluntary wheel running attenuated the decline in sciatic CNAP, while the proteomic profiling identified unique changes in mitochondrial turnover, inflammatory response, and calcium buffering. Similarly, donepezil treatment prevented declines in nerve-stimulated muscle torque production and sciatic CNAP, and the proteomic analysis showed distinct enrichment in mitochondrial immune response, calcium buffering, and inflammatory response pathways. These results underscore the capacity of early interventions to modulate peripheral neurodegenerative processes in 5xFAD mice by targeting distinct proteomic pathways linked to mitochondrial turnover, calcium homeostasis, and inflammation.

The proteomic changes observed at 3, 4, and 7 months in the sciatic nerve of 5xFAD mice share molecular profiles with established AD brain pathology, including alterations in mitochondrial metabolism, calcium homeostasis, and myelination^37,38^. These profiles also overlap with those reported in other neurodegenerative models^39^ and with age-related proteolytic degradation in peripheral tissues^40^, suggesting a conserved neurodegenerative proteomic signature across systems. To our knowledge this is the first study to implicate these proteomic pathways (previously observed only in central AD) in peripheral neuronal tissue. Our findings suggest a broader, systemic characteristic of AD pathology with potential implications for early diagnosis and therapeutic intervention.

While physical activity is recognized as a modifiable risk factor for AD, effectiveness of exercise to reduce AD risk have been inconsistent. For instance, Yu *et al*. reported that six months of cycling training alleviated memory and executive function declines, yet language and attention processing speeds continued to deteriorate. Moreover, the benefits were not sustained over 12 months, as all cognitive measures showed decline^41^. Other studies have indicated that aerobic exercise can benefit those with AD, particularly memory. However, benefits in memory were mostly seen in individuals who had improved their cardiorespiratory fitness, implying that disease stage and may influence adaptive response benefits. Exercise studies using 5xFAD mice have also produced mixed results, some have found that restricting female mice to three hours of running per day improved cognition, neurogenesis in the hippocampus, and reduced amyloid-beta accumulation. In contrast, allowing mice unrestricted access to running wheels did not prevent AD-like pathology, neuroinflammation, or cognitive decline^42^. To build on our previous findings that identified early-onset neuromuscular dysfunction in 5xFAD mice^9^, this study integrated functional and proteomic evidence to reveal a potential molecular basis for these impairments within the peripheral nervous system. Together, our functional and proteomic findings underscore the efficacy of exercise to attenuate of neuromuscular impairment in AD. However, the disease stage for an exercise intervention appears to be critical, as it may be most efficacious when initiated prior to the onset of functional decline. Voluntary wheel running initiated prior to neuromuscular decline preserved sciatic nerve conduction and remodeled the peripheral proteome, enhancing pathways related to mitochondrial quality control, calcium homeostasis, and inflammation regulation. These adaptations are consistent with emerging evidence that exercise promotes neuroprotection through mitochondrial remodeling and immune modulation in both central and peripheral tissues^43–45^. Our findings extend these works by demonstrating that such proteomic remodeling occurs in peripheral nerves in an AD model, reinforcing the concept that early, targeted physical activity may delay or prevent neuromuscular decline through peripheral molecular remodeling.

Although donepezil is recognized as a treatment for cognitive impairment, its effects on neuromuscular function in AD have yet to be fully elucidated. For instance, some studies have indicated that donepezil treatment can improve skeletal muscle mitochondria respiration, and motor function in individuals with MCI and AD^16,18,19^, suggesting potential benefits beyond merely cognitive symptoms, which is in agreement with our present findings. In isolated murine skeletal muscle donepezil treatment prolongs endplate potential and tetanic torque^17^. In mouse models of AD, donepezil administration has shown mixed outcomes in relation to AD pathology in the brain, while very little has been done to explore its effects on neuromuscular function. It is important to note that while donepezil treatment begun prior to neuromuscular impairment attenuated decline, donepezil was unable to rescue neuromuscular dysfunction in 5-month-old 5xFAD mice once impairment had manifested. This temporal benefit of donepezil’s efficacy reflects clinical studies that show declines in physical and cognitive function despite continued treatment^46,47^.

Our study demonstrates that both exercise and donepezil treatment resulted in improvements in neuromuscular function in 5xFAD mice, as evidenced by delayed neuromuscular impairment, enhanced nerve-stimulated muscle torque production, and sciatic nerve proteomic reprofiling. While both interventions modulated shared pathways related to mitochondrial integrity, calcium regulation, and inflammatory response, they also exhibited unique proteomic changes: exercise preferentially enriched pathways associated with metabolic plasticity and mitochondrial turnover, while donepezil modulated cholinergic signaling and immune response elements. These different yet complementary effects suggest a targeted mechanistic foundation for developing new treatments that incorporate the unique benefits of both exercise and donepezil—offering a potentially synergistic approach to preserve peripheral neuromuscular function in AD.

The early manifestation of neuromuscular dysfunction in 5xFAD mice, demonstrates peripheral neuromuscular impairments occur independently or precede central nervous system AD related pathology. Our findings challenge the central dogma that AD pathology is limited to the brain and suggest that peripheral neuromuscular dysfunction precedes cognitive decline through pathways identified in AD brains^37,38^. The two-armed approach of voluntary wheel running and a clinically employed pharmacological intervention (donepezil) in our study offers valuable insights into potential therapeutic strategies for preserving neuromuscular function. However, our study has several limitations. First, the duration of the interventions was relatively short, which may not fully capture the long-term effects of voluntary exercise or donepezil treatment or relate to neuroprotective effects of lifelong exercise^48^. Second, the study focused on a single mouse model (5xFAD), which does not fully represent the complexity of AD pathology in humans. Additionally, this study only focused on male mice. We chose to focus on male 5xFAD mice due to our previous characterization of neuromuscular function impairment being more prominent at 4 months of age in the males^9^. The exclusion of female mice does not discount the need to explore the effects of voluntary wheel running and donepezil in female 5xFAD mice, since they too experience early neuromuscular decline^9^. Integrating lifestyle interventions, such as exercise, with pharmacological treatments could potentially offer multifaceted benefits, addressing both cognitive and neuromuscular health in individuals at risk for, or suffering from AD. By advancing our understanding of the interplay between neuromuscular function and AD pathology, both centrally and peripherally, we can pioneer novel approaches for early interventions and potential treatments for AD.

## Acknowledgements

We thank the Virginia Tech Metabolism Core for technical assistance with skeletal muscle contraction and experiments. We also thank members of JCD’s laboratory for critical feedback and discussion.

## Author Contributions

Study design: MHB, CPN and JCD. Conducting experiments: MHB, DNE, OSW, JB, DB, SH, JKM and CPN. Analyzing data: MHB, JKM, SP, RWG, CPN, and JCD. Manuscript drafting: MHB, OSW, SP, RWG, and JCD.

## Data Availability

The data within this article are available in the article and in its online supplementary material.

---

**Supplemental Figure 1. Proteomic changes in the sciatic nerve in voluntary wheel running:** (A) Left: hierarchical clustering of proteins significantly different between indicated pairings (threshold for clustered proteins was determined by significance between groups). Right: proteins were further grouped using k-means statistics, breaking the significant genes into 10 protein clusters.

**Supplementary Figure 2. STRING network of proteins significantly altered with voluntary wheel running.** Protein-protein interaction networks were generated using STRING analysis (https://string-db.org) to visualize known and predicted associations among significantly abundant proteins in the sciatic nerve of 5xFAD mice following voluntary wheel running. Each node represents a protein, and edges represent functional associations derived from curated databases, experimental data, text mining, co-expression, and gene neighborhood. Edge thickness reflects the confidence of the interaction. Functional enrichment and clustering revealed distinct biological modules associated with exercise-induced proteomic remodeling.

**Supplementary Figure 3. AD-like pathology markers in Voluntary wheel running.** (A) Plasma Neurofilament light chain concentration in WT and 5xFAD. (B) High Molecular weight A-β oligomers in hippocampus. (C) Representative western blot and ponceau n = 6 per WT group and n=7 per 5xFAD Group. Data presented as mean ± SEM and two-way ANOVA was performed and Tukey’s post-hoc text performed when significant interaction between variables was observed (** = p<0.01, *** = p<0.001).

**Supplementary Figure 4. STRING network of proteins significantly altered with donepezil treatment.** STRING analysis (https://string-db.org) was performed on proteins significantly abundant in the sciatic nerve of 5xFAD mice treated with donepezil. Nodes correspond to individual proteins, while edges reflect high-confidence functional interactions based on combined evidence. Functional modules were identified, highlighting pathways responsive to acetylcholinesterase inhibition, including those involved in mitochondrial immune response, inflammation, and calcium handling.

**Supplementary Figure 5. AD-like pathology markers in donepezil treated 5xFAD mice**. (A) Plasma Neurofilament light chain concentration in WT and 5xFAD. (B) High Molecular weight A-β oligomers in hippocampus. (C) Representative western blot and ponceau. n = 6 for WT-Sed, n = 5 for 5xFAD-PLCB group, and n=6 for 5xFAD-DPNZ Group. Data presented as mean ± SEM and two-way ANOVA was performed and Tukey’s post-hoc text performed when significant interaction between variables was observed (**p<0.01, **p<0.01).

**Supplemental Figure 6. Neuromuscular function in 6-month-old 5xFAD mice with donepezil treatment:** (A) Study design of donepezil treatment intervention. (B) Placebo group tibial nerve stimulated, and direct-skeletal muscle stimulated muscle function. (C) Tibial nerve-stimulated skeletal muscle function. (D) Direct muscle stimulated muscle function. (E) In vivo compound nerve action potential (CNAP) of sciatic. n=4 for 5xFAD-PLCB; n= 5 for 5xFAD-DPNZ. Data presented as mean ± SEM and repeated measures and two-way ANOVA was performed and Tukey’s post-hoc test performed when significant interaction between variables was observed (∗P < 0.05, ∗∗P < 0.01). Panel A created in Biorender.

**Supplementary** Figure 7. **STRING network of proteins altered by both voluntary wheel running and donepezil treatment.** A combined STRING network was generated (https://string-db.org) to examine shared and distinct protein-protein interaction patterns among significantly enriched proteins following voluntary wheel running or donepezil treatment. Nodes represent shared or treatment-specific proteins, with edge thickness indicating interaction confidence. Clusters identify biological processes commonly or uniquely influenced by each intervention

**Supplemental Figure 8 . Unique proteomic changes in the sciatic nerve between voluntary and donepezil treatment**: (A) Heat map of proteins uniquely affected by Ex or DPNZ in 5xFAD and maintained to WT-Sed levels. * denotes proteins that group together in STRING clusters. (F) Schematic of protein-protein interacting clusters, representing processes maintained to WT-Sed by either Ex (left) or DPNZ (right) in 5xFAD.

