## Supplementary figures and images for "Age-dependent remodeling of the sciatic proteome in 5xFAD mice can be attenuated by exercise or donepezil treatment to maintain neuromuscular function"

### supplemental figures

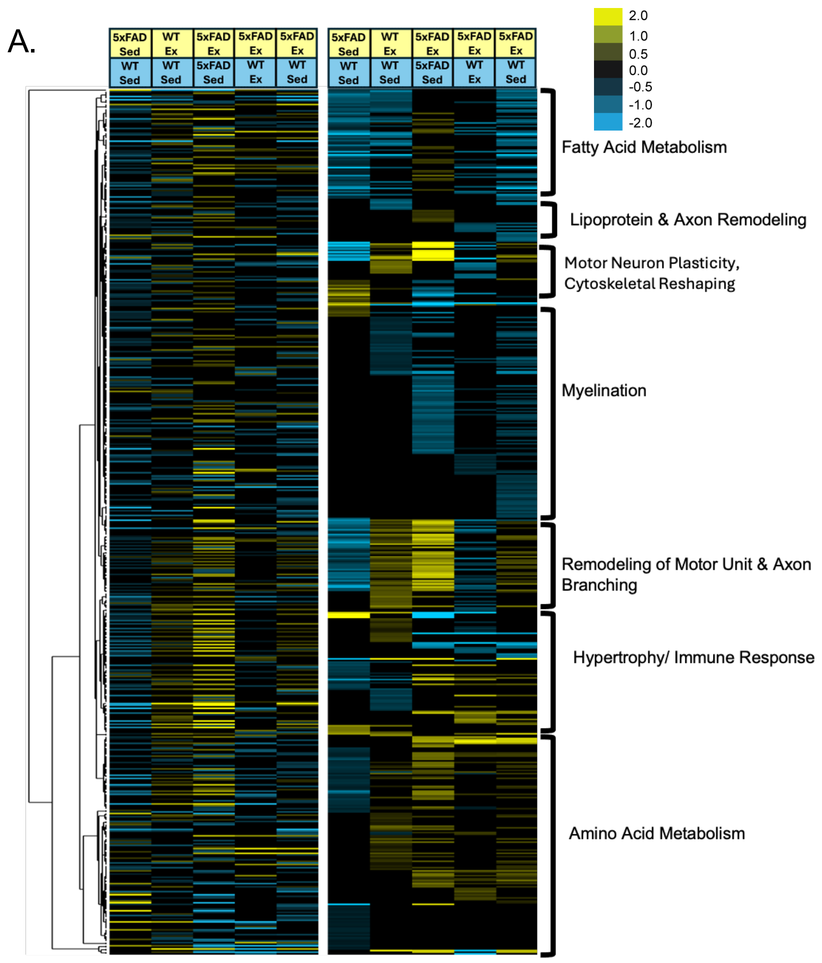

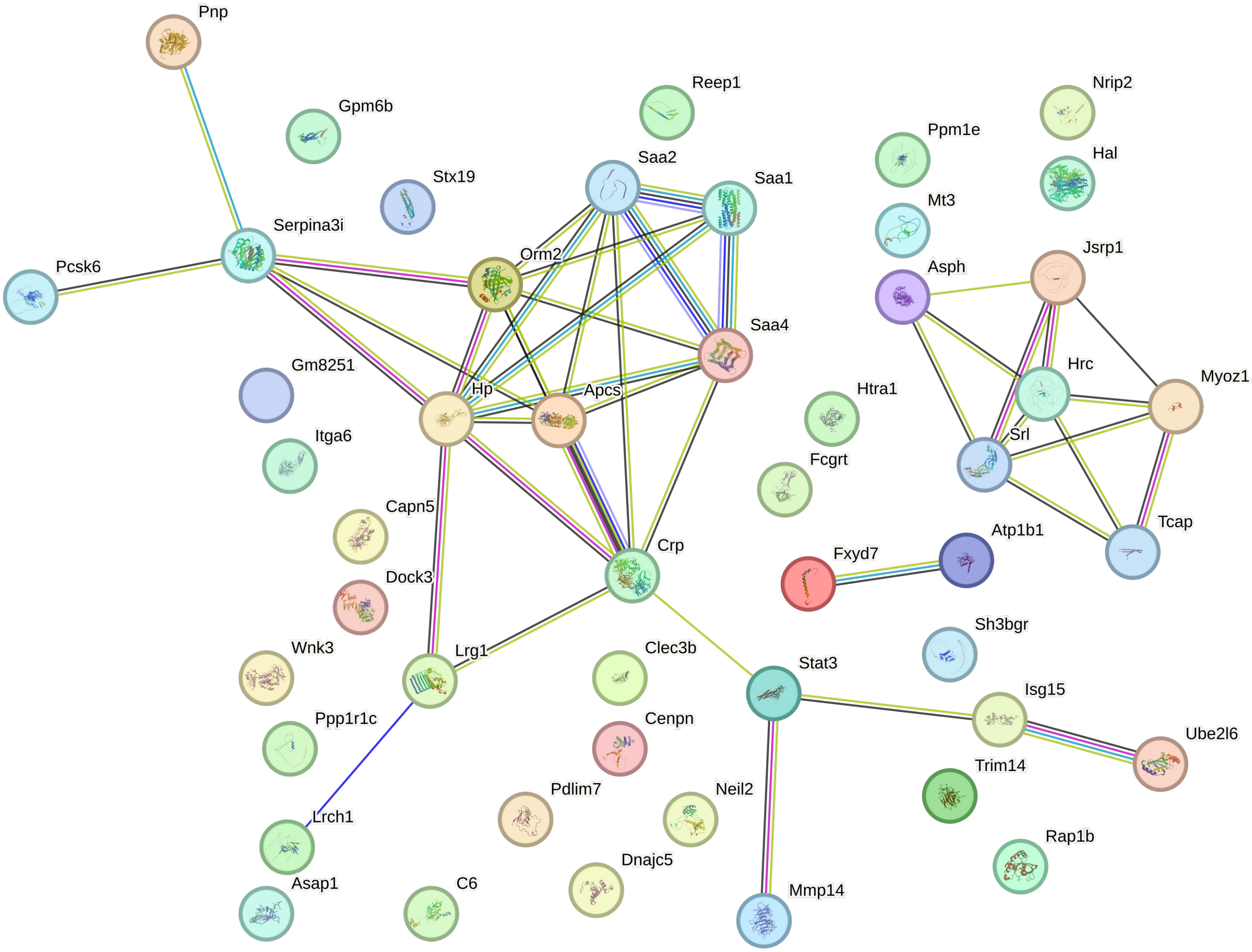

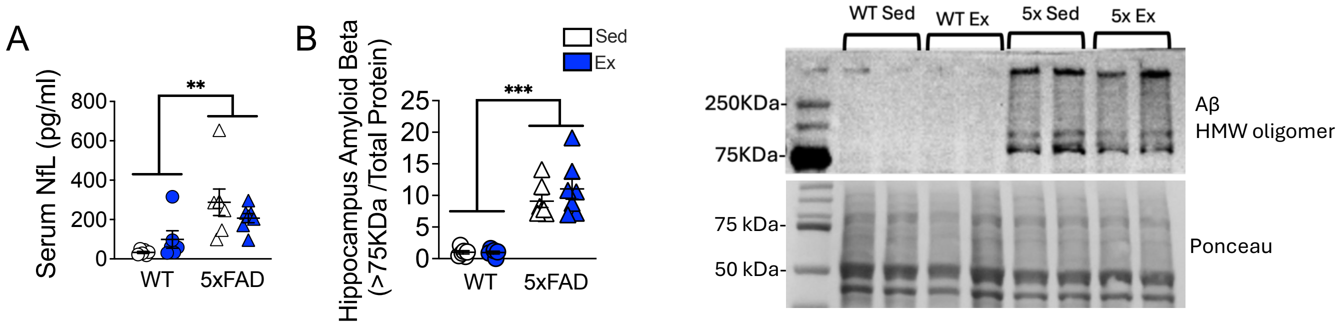

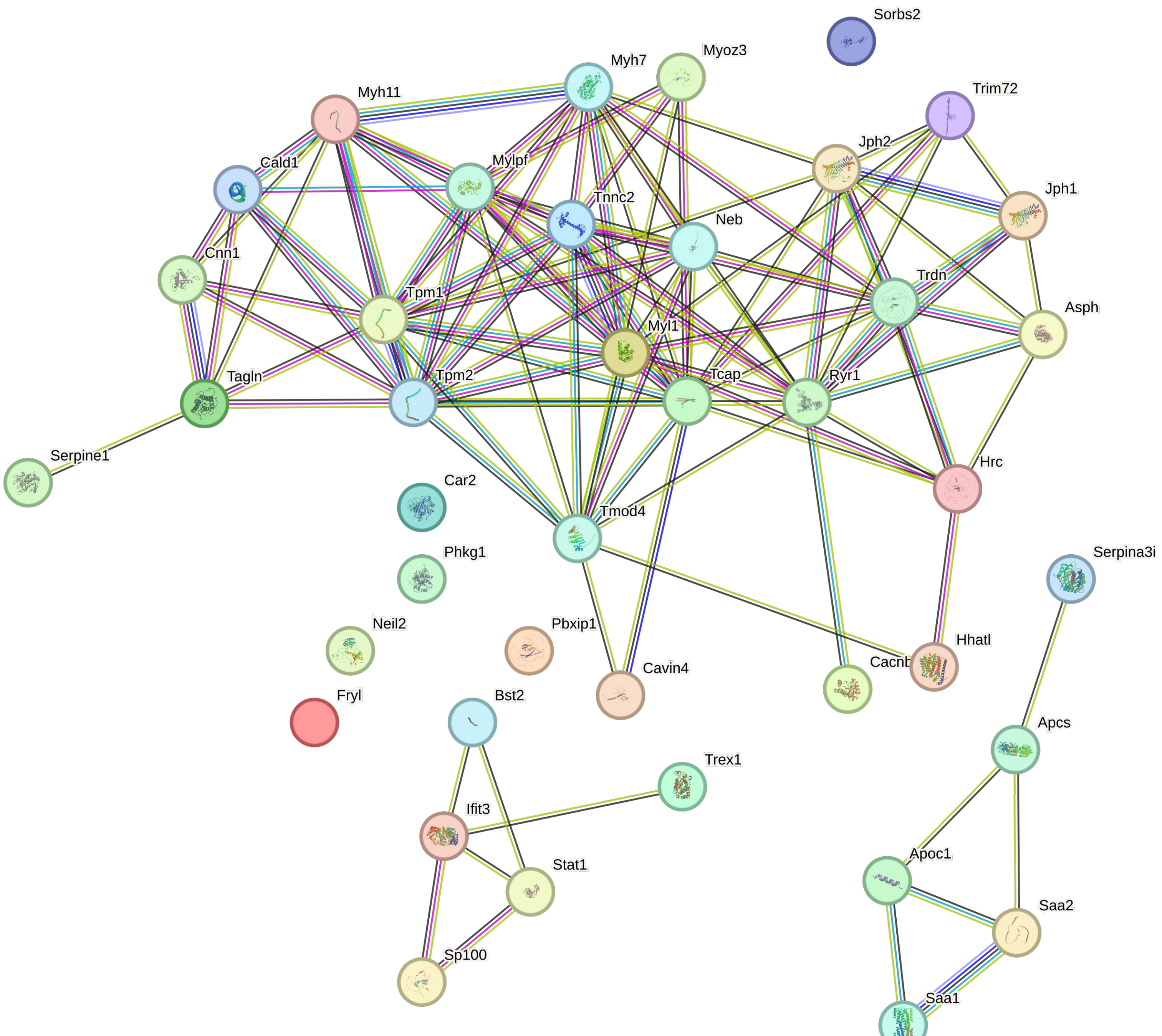

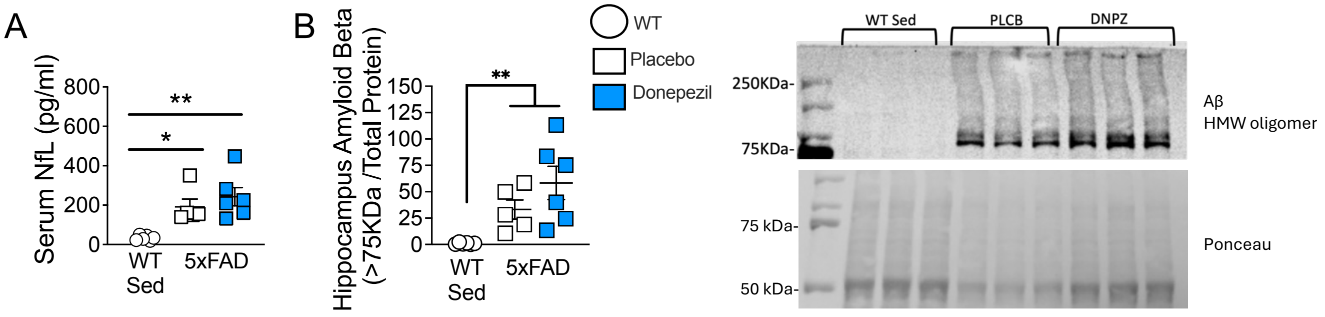

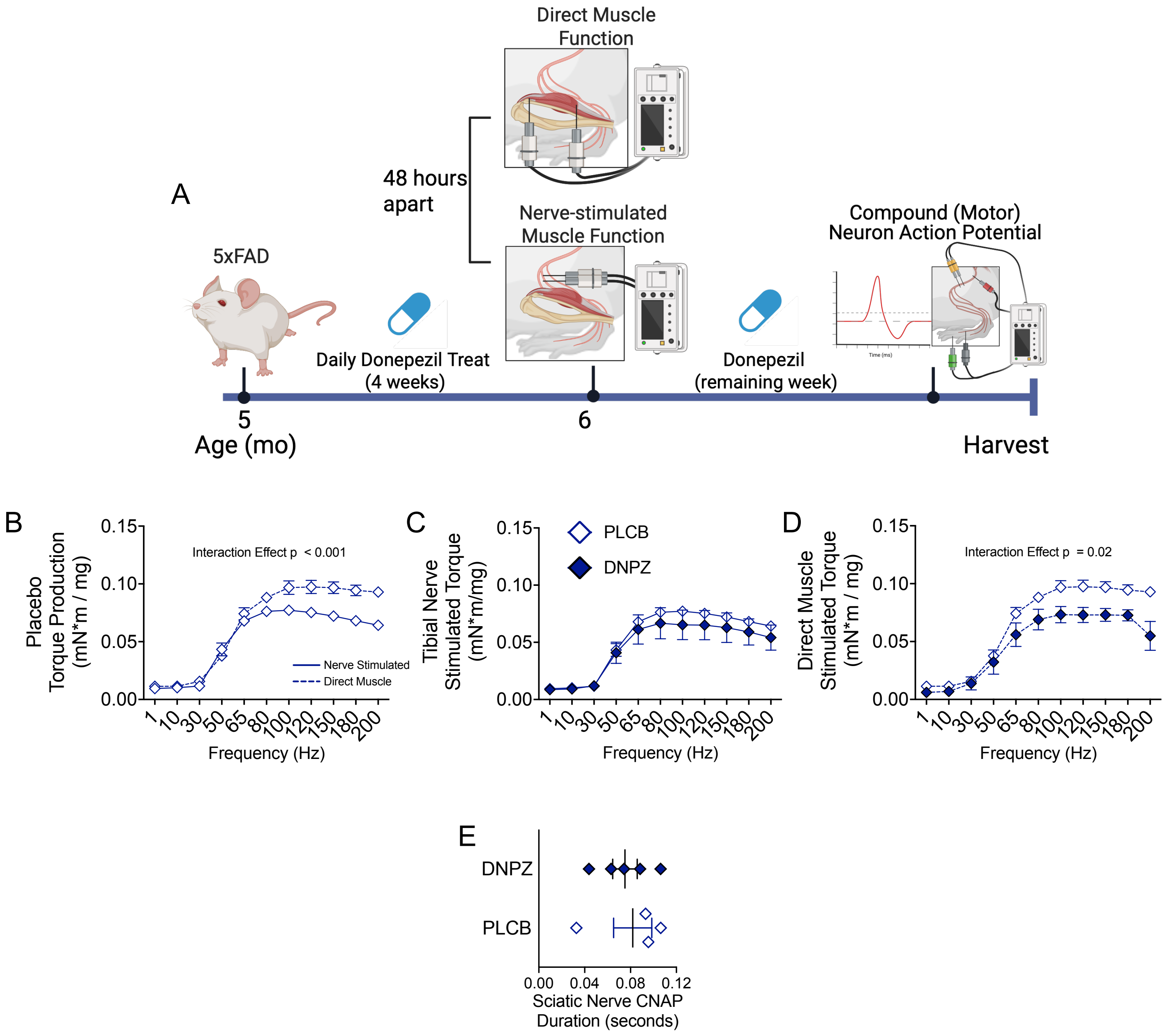

#### Figure S7

Ex

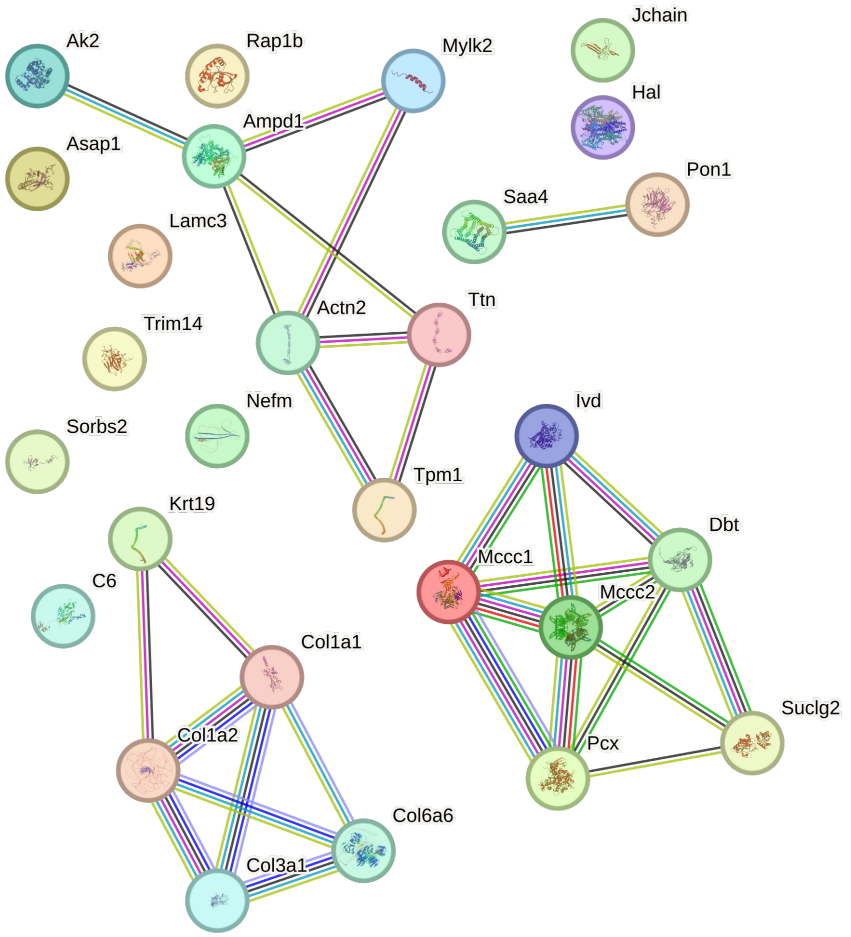


DNPZ

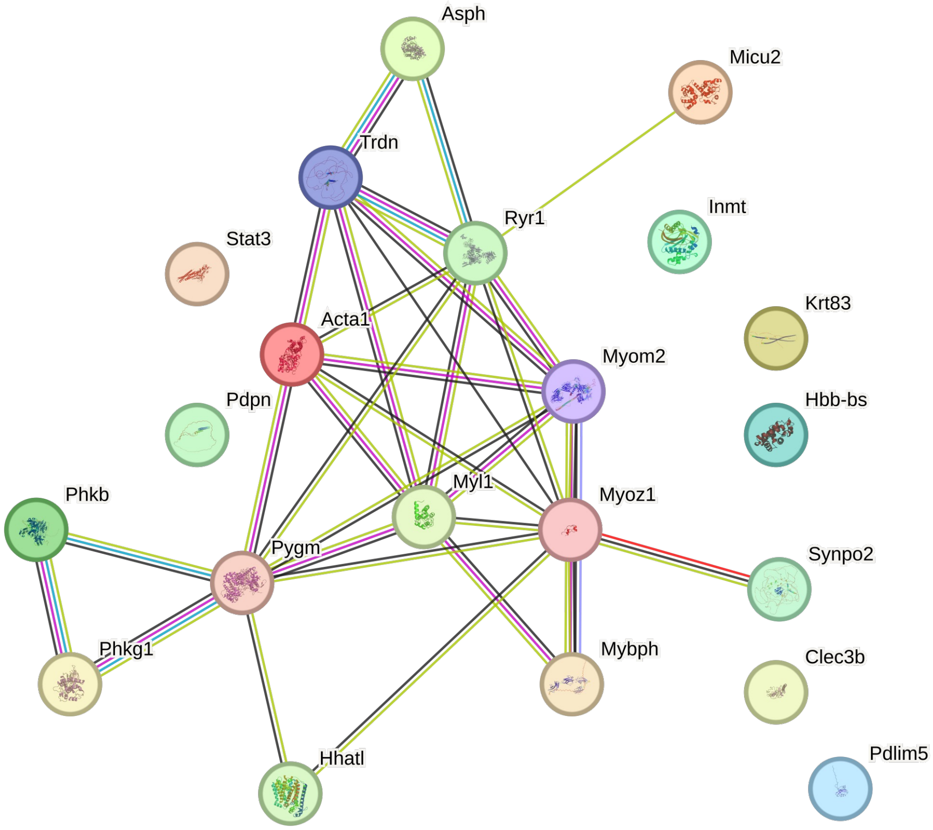

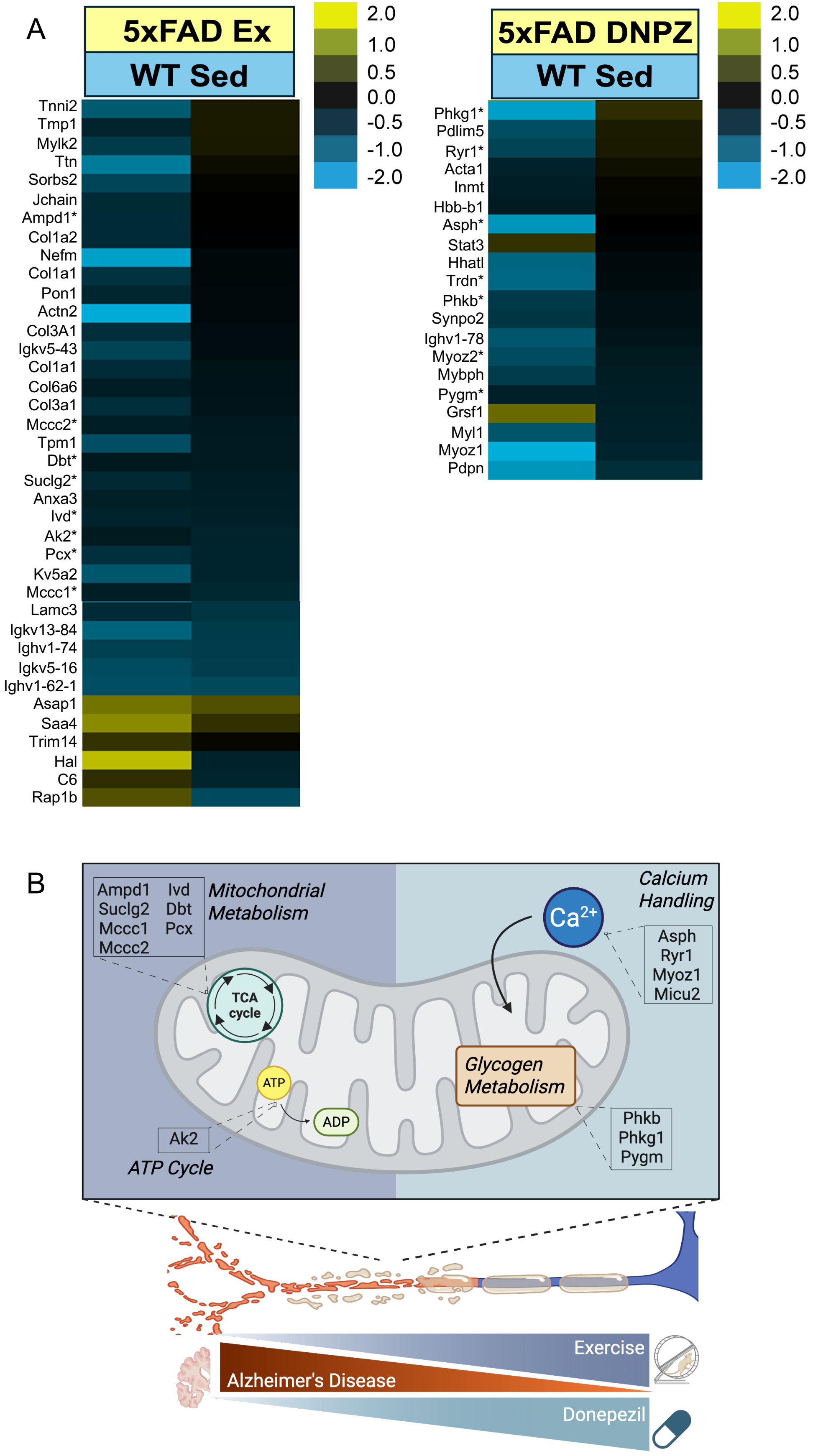
